# COT: an efficient Python tool for detecting marker genes among many subtypes

**DOI:** 10.1101/2021.01.10.426146

**Authors:** Yingzhou Lu, Chiung-Ting Wu, Sarah J. Parker, Lulu Chen, Georgia Saylor, Jennifer E. Van Eyk, David M. Herrington, Yue Wang

## Abstract

We develop an accurate and efficient method to detect marker genes among many subtypes using subtype-enriched expression profiles. We implement a Cosine based One-sample Test (COT) Python software that is easy to use and applicable to multi-omics data. We demonstrate the performance and utility of COT on gene expression and proteomics data acquired from tissue or cell subtypes. Formulated as a one-sample test with Cosine similarity test statistic in scatter space, the detected de novo marker genes will allow biologists to perform a more comprehensive and unbiased molecular characterization, deconvolution and classification of complex tissue or cell subtypes.

## Introduction

Ideally, a molecularly distinct subtype would be composed of molecular features that are uniquely expressed in the cell or tissue subtype of interest while in no others – so called subtype-specific marker genes (SMG) (Wang, Hoffman et al. 2016, Hunt, Freytag et al. 2019). SMG plays a critical role in the characterization, deconvolution, or classification of complex tissue or cell subtypes (Yu, Feng et al. 2010, Parker, Chen et al. 2020). With the increasing availability of subtype-enriched molecular expression profiles acquired by single cell sequencing, cell sorting, microdissection, or mathematical deconvolution, having data-driven software tools to detect SMG, applicable to high throughput multi-omics data, will be essential for facilitating next steps in systems biology research.

We have developed an accurate and efficient method - Cosine based One-sample Test (COT) - to detect SMG among many subtypes using subtype-enriched expression profiles (Fig. 1A). Importantly, COT uses the cosine similarity between a molecule’s cross-subtype expression pattern and the exact mathematical definition of SMG as the test statistic, and formulates the detection problem as a one-sample test. Under the assumption that a significant majority of genes are associated with the null hypothesis, COT approximates the empirical null distribution for calculating p values (Efron 2004). We implement and test the latest functionalities of COT workflow in Python through a three-step assessment. We have previously demonstrated the successful applications of COT prototype on gene expression and proteomics data (Yu, Feng et al. 2010, Chen, Lu et al. 2020).

**Figure 1.**
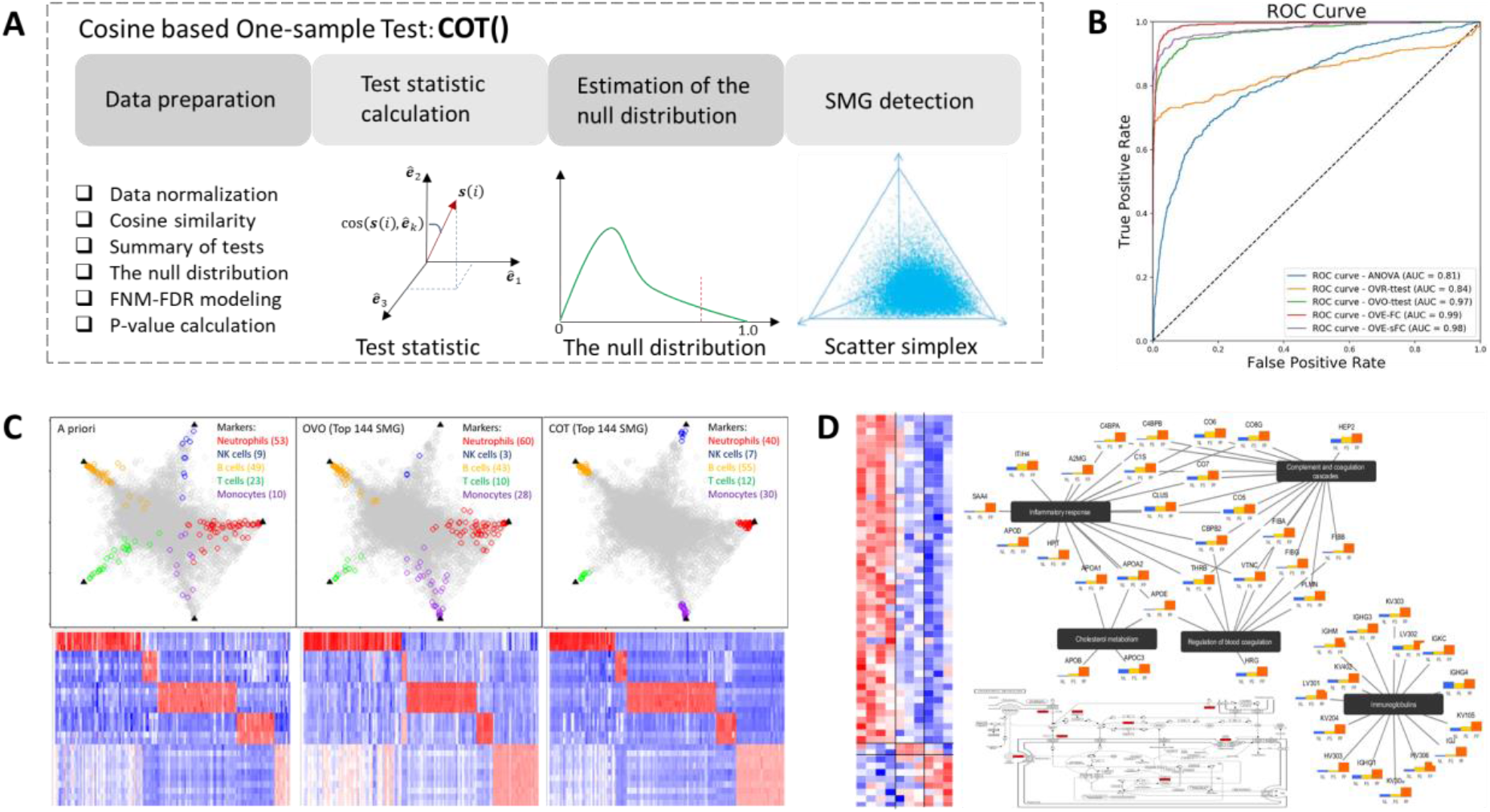
COT workflow and case studies. **A**. Main functional modules in COT Python software. **B**. The receiver operating characteristic curves of four existing methods in overall comparison. **C**. Simplex plots and heatmaps of SMG (color-coded) detected by COT, OVO test, and *a priori* in benchmark assessment (column – protein, row – sample). **D**. Top protein SMG detected by COT on vascular specimens, and top enriched pathways including a detailed cholesterol metabolism circuitry (column – sample, row – protein).

## Results

Mathematically, a SMG of subtype *k* is defined as a gene expressed only in subtype *k*, approximately (Wang, Hoffman et al. 2016, Hunt, Freytag et al. 2019)

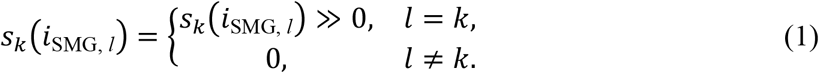

where *s_k_*(*i*_SMG_, *l*) is the expression of gene *i* in subtype *k* in reference to subtype *l*, and is assumed to be nonnegative. Accordingly, the cross-subtype expression patterns of ideal SMG can be concisely represented by the Cartesian unit vectors ***ê**_k_*. Fundamental to the success of the COT method is the newly-proposed test statistic cos(***s***(*i*), ***ê**_k_*) that measures directly the distance between the cross-subtype expression patterns of gene *i* and the ideal SMG of subtype *k* in scatter space. The known cross-subtype expression patterns of ideal SMG associated with the alternative hypothesis permit the use of a one-sample test to detect significant SMG, with 0 < cos(***s***(*i*), ***ê**_k_*) < 1.

In relation to previous work, the effort to detect SMG can be traced back to One-Versus-Rest (OVR) test (Yu, Feng et al. 2010), One-Versus-One (OVO) test (Hunt, Freytag et al. 2019), and most recently One-Versus-Everyone (OVE) test (Wang, Hoffman et al. 2016). We and others have recognized that the test statistics used by most existing methods do not exactly satisfy SMG definition (1) and often require ad hoc OVE set intersection (Yu, Feng et al. 2010, Hunt, Freytag et al. 2019, Chen, Lu et al. 2020).

Following normalization and antilogarithm the COT software tool performs the following major analytics steps (Fig. 1A), (i) Data Preprocessing. Molecule features whose norms of cross-subtype expression levels are lower or higher than pre-fixed thresholds are removed as noise or outliers. (ii) Test Statistic Calculation. For each of remaining genes, cosine similarity cos(***s***(*i*), ***ê**_k_*) between averaged cross-subtype expression patterns and SMG reference is calculated. (iii) Null Distribution Estimation. The empirical null distribution is summarized over all genes and approximated by a mixture of normal distributions. (iv) SMG Detection. Based on the observed test statistic and null distribution, SMG is accepted subject to a proper one-sided significance threshold.

We conduct three-step experiments (overall comparison, benchmark assessment, biomedical case study), on two types of omics data (gene expression and proteomics), to demonstrate the performance of COT Python tool. We first conducted an overall comparison among ANOVA, OVR test, OVO test, and OVE test, using a classical differential analysis setting and simulations generated from real gene expression data (GSE28490). Evaluated by the receiver operating characteristic analyses, this performance survey indicates that OVE/OVO test outperforms most other methods, detailed in Fig. 1B.

We then conducted a benchmark assessment of the COT tool, using a one-sample test setting and the same gene expression data, in comparison with the top performer OVE/OVO test. The geometric proximity of the 144 SMG detected by COT, OVE/OVO, and a priori, to the vertices of scatter simplex and the heatmaps are given in Fig. 1C. The assessment shows that COT tool outperforms the top performer by detecting more ideal SMG.

We further demonstrated the utility of COT tool on experimentally-acquired proteomics data from a cohort of “pure” fibrous plaque (FP), fatty streak (FS), and normal (NL) vascular specimens (Parker, Chen et al. 2020). COT detected 50 FP, 2 FS, and 8 NL markers, respectively. The heatmap and enrichment analysis results are given in Fig. 1D. These proteins are highly consistent with the SMG detected by tissue deconvolution on a much larger cohort (Herrington, Mao et al. 2018). The KEGG maps of FP markers show that nearly all components of the lipoprotein and immunoglobulin pathways have been detected. Intriguingly, two FS markers have previously been reported functionally involved in vascular pathology and atherosclerosis progresses.

## Discussion and outlook

The COT software tool provides an accurate data-driven marker detection tool where the test statistic matches exactly the definition of SMG and permits the novel formulation of a one-sample test. Furthermore, the test statistic cos(***s***(*i*), ***ê**_k_*) is calculated in reference to the alternative hypothesis, making the resulting p-values more meaningful and avoiding the caveat in general significant tests with no knowledge on the alternative hypothesis (Efron 2004). Moreover, COT is efficient in that neither OVE set intersection nor intractable sample permutation is needed (Hunt, Freytag et al. 2019, Chen, Lu et al. 2020). While the case study here involves only transcripts and proteins, COT tool is applicable to other omics types (Supplementary Information). The nonnegativity requirement in COT is critical as similarly imposed by classical t-ratio test.

Fundamental to the success of the COT method is the newly-proposed test statistic cos(***s***(*i*), ***ê**_k_*) that measures directly the distance between the cross-subtype expression patterns of gene *i* and the ideal SMG of subtype *k* in scatter space. The known cross-subtype expression patterns of ideal SMG associated with the alternative hypothesis permit the use of a one-sample test to detect significant SMG, making the resulting p-values more meaningful in the framework of Bayes hypothesis testing and avoiding the caveat in general significant tests with no knowledge on the alternative hypothesis (Efron 2004).

More importantly, COT framework is efficient in that neither OVE set intersection nor intractable sample permutation is needed in estimating the null distribution (Hunt, Freytag et al. 2019, Chen, Lu et al. 2020), where a significant majority of genes are assumed to be associated with the null hypothesis. The null distribution plays a crucial role in large-scale multiple testing. However, because the size of subtype-enriched samples is often relatively small and the non-SMG patterns are highly complex, classical methods to estimate the null distribution in a two-sample test setting is impractical or intractable (Chen, Lu et al. 2020) and in appropriate (Efron 2004). The important assumption is that due to the significant large proportion of null features, the data can show the null distribution itself (Efron 2004). The theoretical null may fail in some cases, which is not completely wrong but needs adjustment accordingly. In our study, we adopted a mixture of five normal distributions (finite normal mixture – FNM) to approximate the empirical COT distribution (Equihua 1988, Wang, Adali et al. 1997). Moreover, instead of modeling only the null distribution and using FDR-guided iterative estimation strategy, one can also simultaneously model and estimate both the null and alternative distributions using a proper mixture distribution when the number of features associated with the alternative hypothesis is sufficient (Wang, Adaly et al. 1998).

In relation to our own recent work (Chen, Lu et al. 2020), the so-called subtype-specific differentially-expressed genes (SDEG) are defined as being most-upregulated in only one subtype but not in any other. In other words, we considered any case with at least ‘two-equal winners’ as non-SDEG. Clearly, SMG is different from and much more stringent than SDEG.

In large-scale multiple testing, standardization of test statistic to summarize a unified null distribution is widely adopted for two reasons, i.e., information cross-fertilization and comparable z-value conversion among all genes. In classical methods, the test statistics are often scaled by estimated sampling standard deviation using either parametric formula or bootstrapping. In our work, importantly, COT uses a cosine score function that is norm/magnitude invariant so that the variance stabilization is automatically achieved.

## Methods

An important but frequently underappreciated issue is how best to define and detect a cell or tissue subtype-specific marker genes (SMG) among many subtypes. Ideally, a molecularly distinct subtype would be composed of molecular features that are uniquely expressed in the cell or tissue subtype of interest while in no others – so called subtype-specific marker genes (SMG) (Kuhn, Thu et al. 2011, Hart, Sheftel et al. 2015, Newman, Liu et al. 2015, Chen, Wu et al. 2020, Patrick, Taga et al. 2020).

The most frequently used methods rely on an ANOVA model where the null hypothesis states that samples in all subtypes are drawn from the same population. Another population method is the One-Versus-Rest Fold Change test or One-Versus-Rest t-test (OVR-FC/t-test) that is based on the ratio of the average expression in a particular subtype to the averaged expression in all other (rest) samples (Shoemaker, Lopes et al. 2012, Zhang, Chen et al. 2014, Chikina, Zaslavsky et al. 2015). Alternative strategies include One-Versus-One (OVO) t-test and Multiple Comparisons with the Best (MCB) (Hsu 1996, Wang, Master et al. 2006, Newman, Liu et al. 2015). Importantly, we and others have recognized that the test statistics used by most existing methods do not exactly satisfy SMG definition and often ad hoc OVE set intersections have been used to finalize SMG (Wang, Hoffman et al. 2016, Chen, Lu et al. 2020, Patrick, Taga et al. 2020).

To address the critical problem of the absence of accurate SMG detection methods, we have previously proposed, tested and applied One-Versus-Everyone Fold Change (OVE-FC) test (Yu, Feng et al. 2010, Yu, Li et al. 2011) and more recently blended OVE/OVO permutation/t-test (Chen, Lu et al. 2020). We have demonstrated real biomedical utilities of these COT prototypes on gene expression and proteomics data for the purpose of characterizing or classifying complex subtypes. These applications have led to novel findings and hypotheses (Herrington, Mao et al. 2018, Parker, Chen et al. 2020). The COT software tool reported here evolves from our previous work, while uses the cosine similarity between a molecule’s cross-subtype expression pattern and the exact mathematical definition of SMG as the test statistic, and formulates the detection problem as a one-sample test (Efron 2004). The COT Python package implemented and tested the latest functionalities of the COT workflow.

Based on the SMG definition, COT uses the cosine similarity between a molecule’s cross-subtype expression pattern and the exact mathematical definition of SMG as the test statistic, and formulates the detection problem as a one-sample test. Specifically, for gene *i* and subtype *k*, the COT test statistic is given by

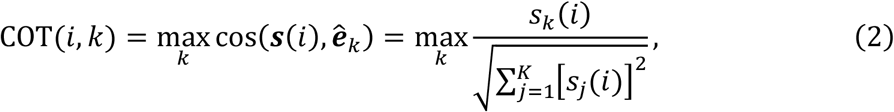

where ***s***(*i*) = [*s*_1_(*i*), *s*_2_(*i*),…, *s_K_*(*i*)] is the averaged cross-subtype expression pattern of gene *I* over samples, ***ê**_k_* is the Cartesian unit vector representing SMG definition of subtype *k*, and *K* is the number of subtypes. Because ***s***(*i*) is confined within the first quadrant whose central vector is the all-ones vector 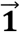, we have 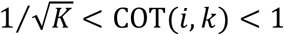. Note that the ‘argmax’ operation in (2) is specifically applied to satisfy the dichotomized setting of the null and alternative hypotheses, considering that the alternative hypothesis is intrinsically associated with multiple subtypes.

We used a mixture of *K* normal distributions (finite normal mixture – FNM model) to approximate the empirical COT distribution (Equihua 1988, Efron 2004). The FNM distribution is initialized by agglomerative clustering and then estimated by the expectation-maximization (EM) algorithm (Wang, Adali et al. 1997, Wang, Adaly et al. 1998). We then improved the approximation by an FDR-guided iterative estimation strategy, where the same number of ‘falsely accepted’ SMG, as predicted by the q-value, are randomly/uniformly removed from the next iteration of the approximation (Storey and Tibshirani 2003, Efron 2004). This procedure converges to a stationary point usually within 5~20 iterations with adjusted collective cut-off p-values of 0.001, 0.005, 0.01, and 0.05.

Alternatively, a molecularly distinct subtype may be characterized by molecular features that are uniquely silent in the cell or tissue subtype of interest while in no others – so called subtype-specific downregulated genes (SDG) (Yu, Feng et al. 2010). Mathematically, a SDG of subtype *k* is defined as a gene expressed only in subtype *k*, approximately

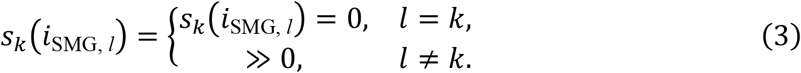

where *s_k_*(*i*_SMG_, *l*) is the expression of gene *i* in subtype *k* in reference to subtype *l*, and is assumed to be nonnegative. A modified COT test statistic can be used to detect SDG given by

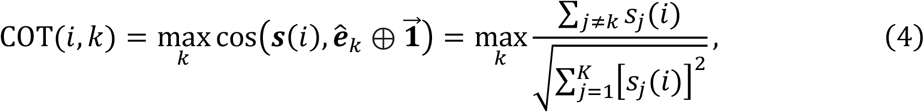

where ⊕ is the exclusive disjunction XOR operation (Yu, Feng et al. 2010).

We implement and test the latest functionalities of COT workflow in Python. The Python module is free online at GitHub (https://github.com/MintaYLu/COT), which was built on top of NumPy and Pandas, and distributed under the MIT license.

The rows of input data matrix should correspond to genes, and the columns correspond to samples. Note that the subtype label is needed for each sample for COT test statistics.

### Data preprocessing

Normalization and batch effect removal are conventional procedures to reduce technical or experimental variations in molecular expression profiles prior to any further analysis. Many normalization pipelines or packages have been developed to reduce technical variations in a specific molecular measurement platform, *e.g*., RMA and PLIER for Affymetrix Microarray, SWAN and BMIQ for Infinium HumanMethylation450 array. Most of the public dataset repositories provide normalized data and the description of normalization workflow. Batch effect is the systematic error introduced by the time- and site-dependent experimental variations. ComBat (Li, Johnson et al. 2006) is a popular and effective method to remove batch effects.

## Supplementary results

### Overall comparison studies

In the overall comparative studies among ANOVA, OVR test, OVO test, and OVE test (Chen, Lu et al. 2020), we used a classical differential analysis setting and simulations generated from real gene expression data (GSE28490). Specifically, simulation datasets are generated by assigning a portion of the genes as being most-upregulated in only one subtype but not in any other, with fold change drawn in certain ranges. To recapitulate the characteristics of real expression data, we used the parameter values estimated from real data. We used both ROC curves and partial area under ROC curve (pAUC) to evaluate the performances of the selected existing methods. Evaluated by the receiver operating characteristic analyses, this performance survey indicates that OVE/OVO test outperforms most other methods, see more details in Figure 1B and also in our recently published report (Chen, Lu et al. 2020).

### Experimental assessment of COT method on benchmark datasets

We then conducted a benchmark assessment on COT tool (Figure 1C), using a one-sample test setting and the real gene expression data (GSE28490). Figure S2 shows the empirical distribution (histogram) of COT test statistic obtained from the dataset, where the lower bound matches well the expected value of 0.447. The initial fitting is given in Figure S3. Figure S4 shows the converged FNM approximation of COT distribution after about 6 iterations. Figure S5 shows the empirical distribution of COT p-value calculated using the converged FNM approximation of COT distribution. Table S1 provides the corresponding p-value threshold, q-value, COT threshold, and number of accepted SMG associated with the converged FNM approximation of COT distribution.

### Application of the COT tool on proteomics data of vascular specimens

We applied the COT software tool to detect SMG on two independently acquired proteomics mass spectrometry datasets. The first dataset is acquired from a cohort (n=10) of “pure” fibrous plaque (FP), fatty streak (FS), and normal (NL) vascular specimens (Parker, et al., 2020). The second dataset is acquired from a cohort (n=78) of heterogeneous vascular specimens containing mixed FP, FS and NL subtypes (Herrington, Mao et al. 2018). The experimental results show that the detected two sets of SMGs are highly consistent across these two datasets.

Table S2 provides a list of the top 60 SMG detected by COT on proteomics data of pure samples. The analysis on the ‘pure’ specimen dataset detected proteins enriched in most of the same pathways we have seen previously (Parker, Chen et al. 2020), including many of the immunoglobulins (indicative of immune activation and B cell involvement), complement factors (indicative of immune activity and inflammation), as well as protein degradation, and many of the apolipoproteins which shuttle cholesterols from the liver to the periphery and back to the liver again. Some preliminary evidence at the mRNA level show that at least the transcripts of apolipoproteins are present in vasculature of the aorta or LAD, indicating that perhaps they are truly being translated from DNA and transcribed locally. The two KEGG maps (Figures S6 and S7) show that nearly all components of the lipoprotein molecules have been detected. The Complement figure shows that proteins involved in the later phase of complement activation seem to dominate the FP markers, which may have some functional significance. The Cytoscape map puts them all in perspective (Figure S8).

The fatty streaks FS markers are intriguing. GBB2 is a fairly generic signaling molecular that acts in concert with G-protein receptors, but Angiotensin II is one of those receptors that is known to be pretty involved in vascular pathology, as is beta adrenergic receptor which is also quite a potent actor in vascular physiology. Then there is UGDH - which is an enzyme involved in the synthesis of glycosaminoglycans (GAGs). It has been reported (Stevens, Colombo et al. 1976) that an overall decrease in GAGs as atherosclerosis progresses, and early stage lesions (i.e., fatty streaks) have a transient spike in some GAGs that then regresses as the lesion gets worse. The UDGH enzyme is kind of a rate limiting, first step enzyme in this process and its upregulation can drive probably either selective or generic GAG synthesis (Clarkin, Allen et al. 2011).

The normal marker NL proteins are an interesting selection. K2C8 is a cytoskeletal keratin involved in the contractile apparatus of striated muscle and may also play a similar role in linking contractile apparatus of SMC to the desmosome (thus, involved in normal SMC function). Along those lines MYL9 is also involved in SMC contraction, and LIMS1 is important for cell-cell adhesion and cell survival via integrin signaling, so probably also an indicator that SMCs are linking up in their normal way to each other. The others are involved in general maintenance functions like ER folding and ribosome. More specifically, the normal markers of contractile apparatus, adhesion, and ECM include K2C8, MYL9, LIMS1, SPRL1, SPON1, and of protein synthesis, folding and quality control include RL11, HSP74, DJC10. Figure S9 shows the geometric proximity of top SMG.

The objective of detecting SMG using the second dataset acquired from ‘heterogeneous’ rather than ‘pure’ specimens is to cross-validate the SMG detected from a small ‘pure’ sample cohort using an independent and much larger ‘heterogeneous’ sample cohort. We first applied state-of-the-art unsupervised deconvolution tool, Convex Analysis of Mixtures (CAM) (Wang, Hoffman et al. 2016, Chen, Wu et al. 2020), to identify the ‘transformed’ reference of ideal SMG in the scatter space, i.e., the vertices of the scatter simplex. We then conducted COT in the scatter simplex of heterogeneous specimen dataset. Table S3 shows the consistency between the SMGs detected from the pure and heterogeneous samples. Note that FS is considered a ‘transitional’ subtype between NL and FP, therefore the ‘cross-talk’ of FS markers with NL and FP is expected (Herrington, Mao et al. 2018, Parker, Chen et al. 2020).

**Table S1.** The corresponding p-value threshold, q-value, COT threshold, and number of accepted SMG associated with the converged FNM approximation of COT distribution in dataset GSE28490 using an FDR-guided iterative estimation strategy.

| P-value | Q-value | COT | # of significant |
| --- | --- | --- | --- |
| 1 | 1 | 0.45 | 20367 |
| 0.05 | 0.58 | 0.87 | 1750 |
| 0.01 | 0.25 | 0.95 | 829 |
| 0.005 | 0.16 | 0.96 | 639 |
| 0.001 | 0.060 | 0.987 | 336 |
| 0.0007 | 0.050 | 0.9905 | 283 |
| 0.0001 | 0.020 | 0.9984 | 100 |

**Table S2.** List of top 60 SMG detected by COT on proteomics data of pure samples.

| Uniport | Gene | MaxCos | Subtype |
| --- | --- | --- | --- |
| P01023 | A2MG | 0.984953 | FP |
| P35858 | ALS | 0.949705 | FP |
| P04114 | APOB | 0.932073 | FP |
| P05090 | APOD | 0.916104 | FP |
| P02649 | APOE | 0.940183 | FP |
| P04003 | C4BPA | 0.941963 | FP |
| P20851 | C4BPB | 0.889659 | FP |
| Q96IY4 | CBPB2 | 0.874947 | FP |
| P02671 | FIBA | 0.938695 | FP |
| P02675 | FIBB | 0.962863 | FP |
| P02679 | FIBG | 0.966014 | FP |
| P00738 | HPT | 0.973683 | FP |
| P01871 | IGHM | 0.932454 | FP |
| P01591 | IGJ | 0.987092 | FP |
| P19823 | ITIH2 | 0.926847 | FP |
| Q96PD5 | PGRP2 | 0.886458 | FP |
| P00747 | PLMN | 0.863939 | FP |
| P55058 | PLTP | 0.977319 | FP |
| O00391 | QSOX1 | 0.915904 | FP |
| P35542 | SAA4 | 0.967419 | FP |
| P04004 | VTNC | 0.926702 | FP |
| O60701 | UGDH | 0.843959 | FS |
| P00450 | CERU | 0.855635 | FP |
| P00734 | THRB | 0.862961 | FP |
| P01031 | CO5 | 0.841316 | FP |
| P01597 | KV105 | 0.879806 | FP |
| P01617 | KV204 | 0.911765 | FP |
| P01621 | KV303 | 0.891841 | FP |
| P01625 | KV402 | 0.912694 | FP |
| P01714 | LV301 | 0.916399 | FP |
| P01764 | HV303 | 0.965693 | FP |
| P01767 | HV306 | 0.900022 | FP |
| P01834 | IGKC | 0.887244 | FP |
| P01857 | IGHG1 | 0.859667 | FP |
| P01860 | IGHG3 | 0.888342 | FP |
| P01861 | IGHG4 | 0.905672 | FP |
| P01876 | IGHA1 | 0.938344 | FP |
| P02647 | APOA1 | 0.934599 | FP |
| P02652 | APOA2 | 0.90552 | FP |
| P02656 | APOC3 | 0.916256 | FP |
| P04196 | HRG | 0.854048 | FP |
| P04433 | KV309 | 0.956962 | FP |
| P05546 | HEP2 | 0.840869 | FP |
| P05787 | K2C8 | 0.888285 | NL |
| P07360 | CO8G | 0.874533 | FP |
| P08185 | CBG | 0.840799 | FP |
| P09871 | C1S | 0.849535 | FP |
| P10643 | CO7 | 0.890565 | FP |
| P10909 | CLUS | 0.84181 | FP |
| P13671 | CO6 | 0.861453 | FP |
| P19827 | ITIH1 | 0.861765 | FP |
| P24844 | MYL9 | 0.848724 | NL |
| P34932 | HSP74 | 0.843921 | NL |
| P48059 | LIMS1 | 0.8543 | NL |
| P62879 | GBB2 | 0.842642 | FS |
| P62913 | RL11 | 0.846606 | NL |
| P80748 | LV302 | 0.879469 | FP |
| Q14515 | SPRL1 | 0.848788 | NL |
| Q14624 | ITIH4 | 0.863949 | FP |
| Q9HCB6 | SPON1 | 0.90266 | NL |

**Table S3.** Overlap between the top SMGs detected from purified and “bulk” samples.

| SMG of heterogeneous samples | SMG of pure sample cohort |  |  |  |
| --- | --- | --- | --- | --- |
|  | Top 60 | FP(51) | FS(2) | NL(7) |
|  | S1 | 42 | 0 | 0 |
|  | S2 | 9 | 2 | 1 |
|  | S3 | 0 | 0 | 6 |

**Figure S1.**
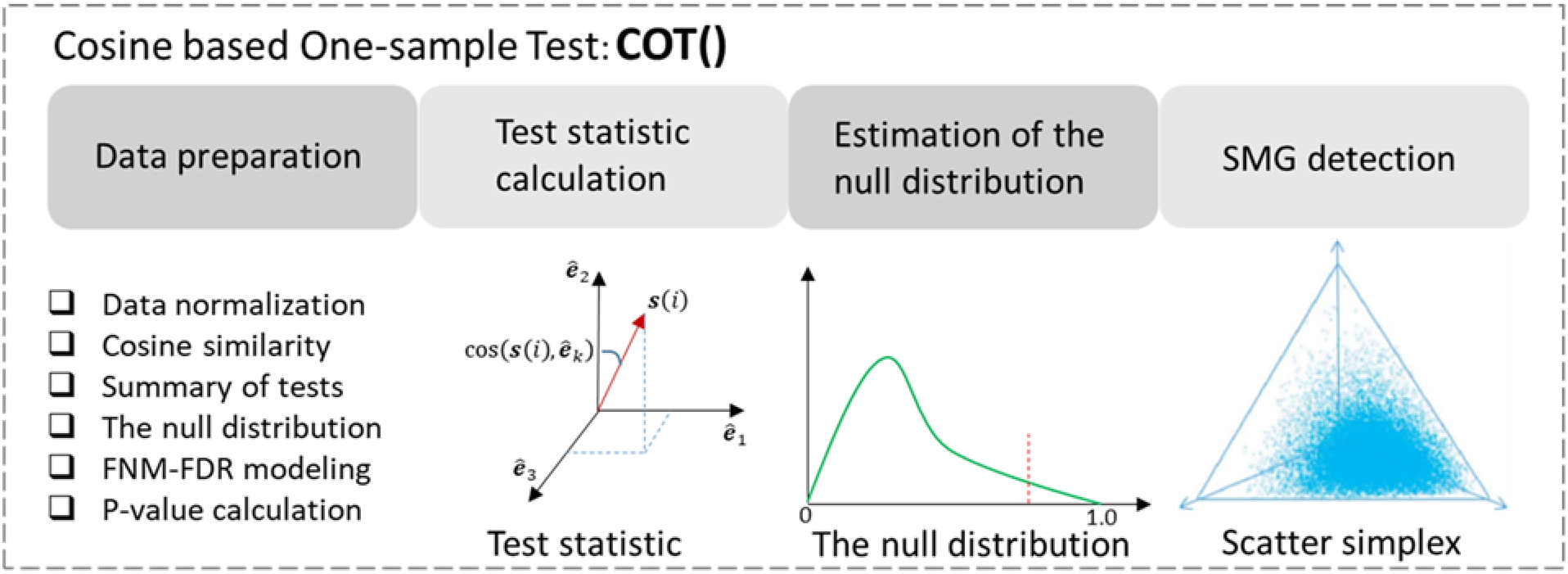
The overall workflow diagram of the COT Python package.

**Figure S2.**
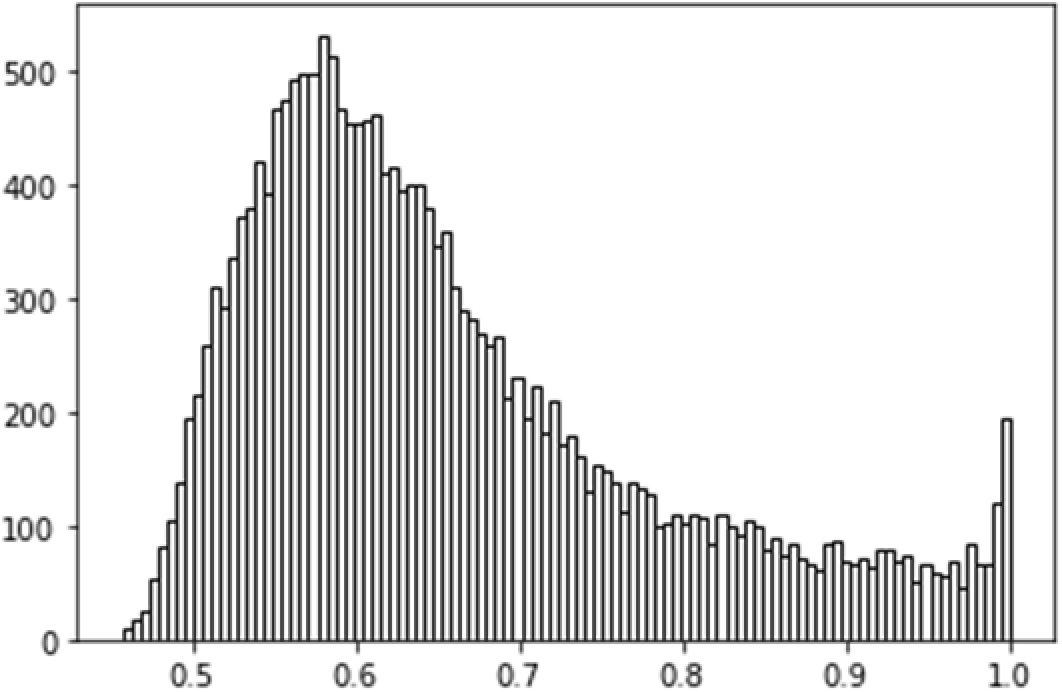
The distribution (histogram) of COT test statistic in dataset GSE28490.

**Figure S3.**
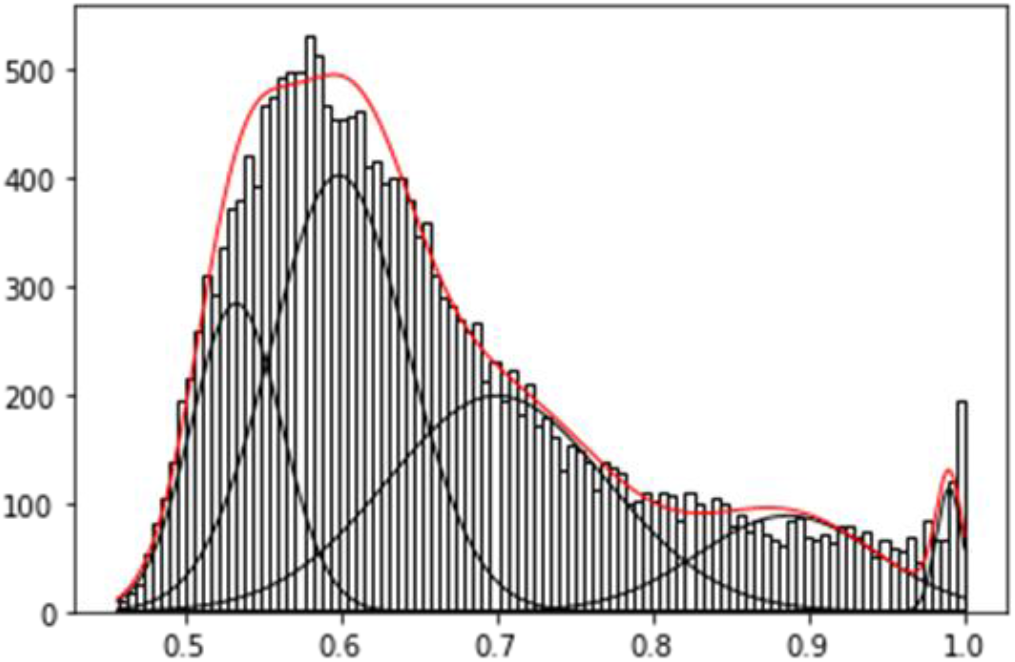
The initial fitting of COT distribution in dataset GSE28490 using a mixture of five normal distributions (the black curves are the estimated individual Gaussian kernels and the red curve is the overall mixture distribution), without FDR-guided adjustment.

**Figure S4.**
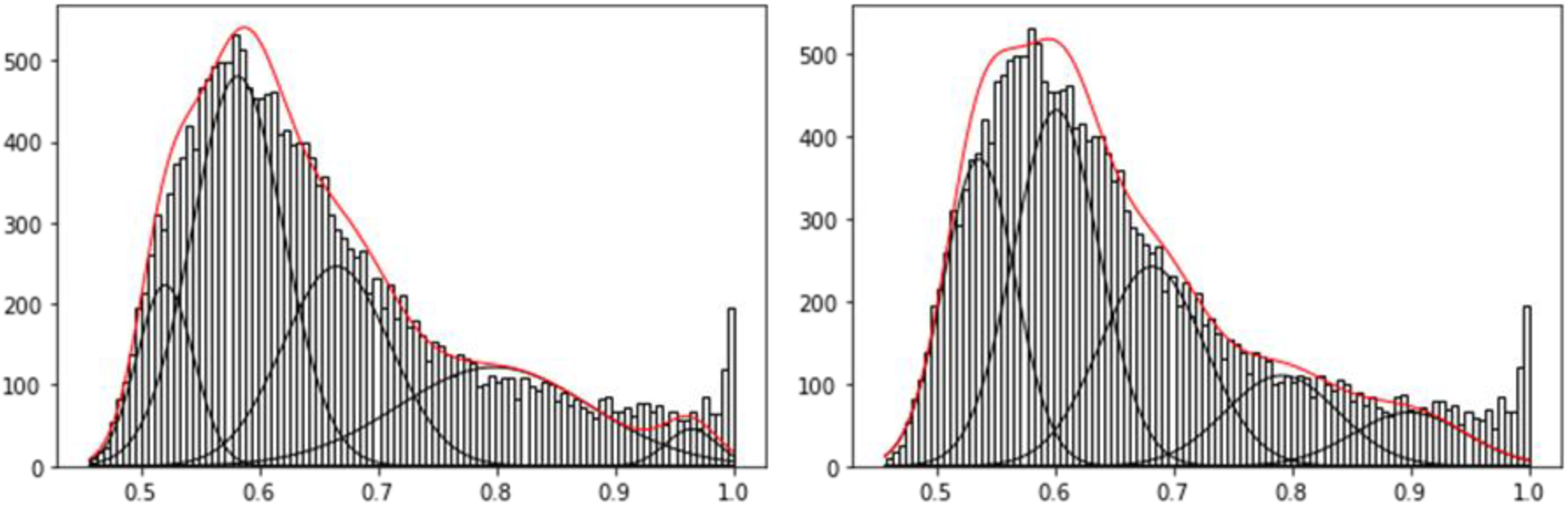
The converged FNM approximation of COT distribution in dataset GSE28490 using an FDR-guided iterative estimation strategy (left: 4 iterations, right: 6 iterations).

**Figure S5.**
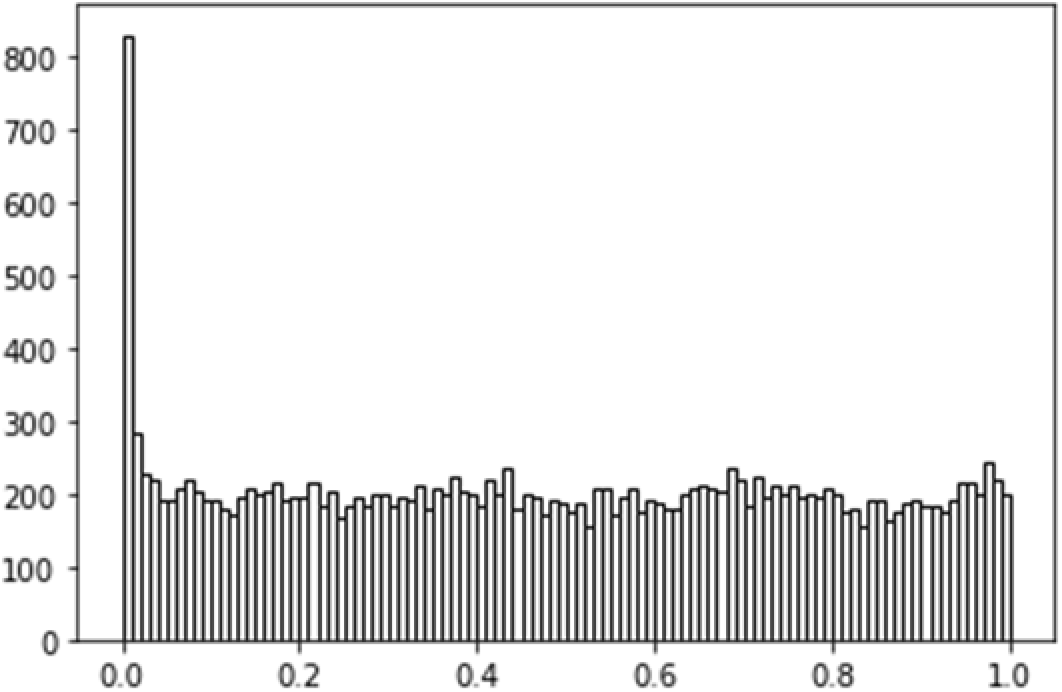
The empirical distribution of COT p-value calculated using the converged FNM approximation of COT distribution in dataset GSE28490.

**Figure S6.**
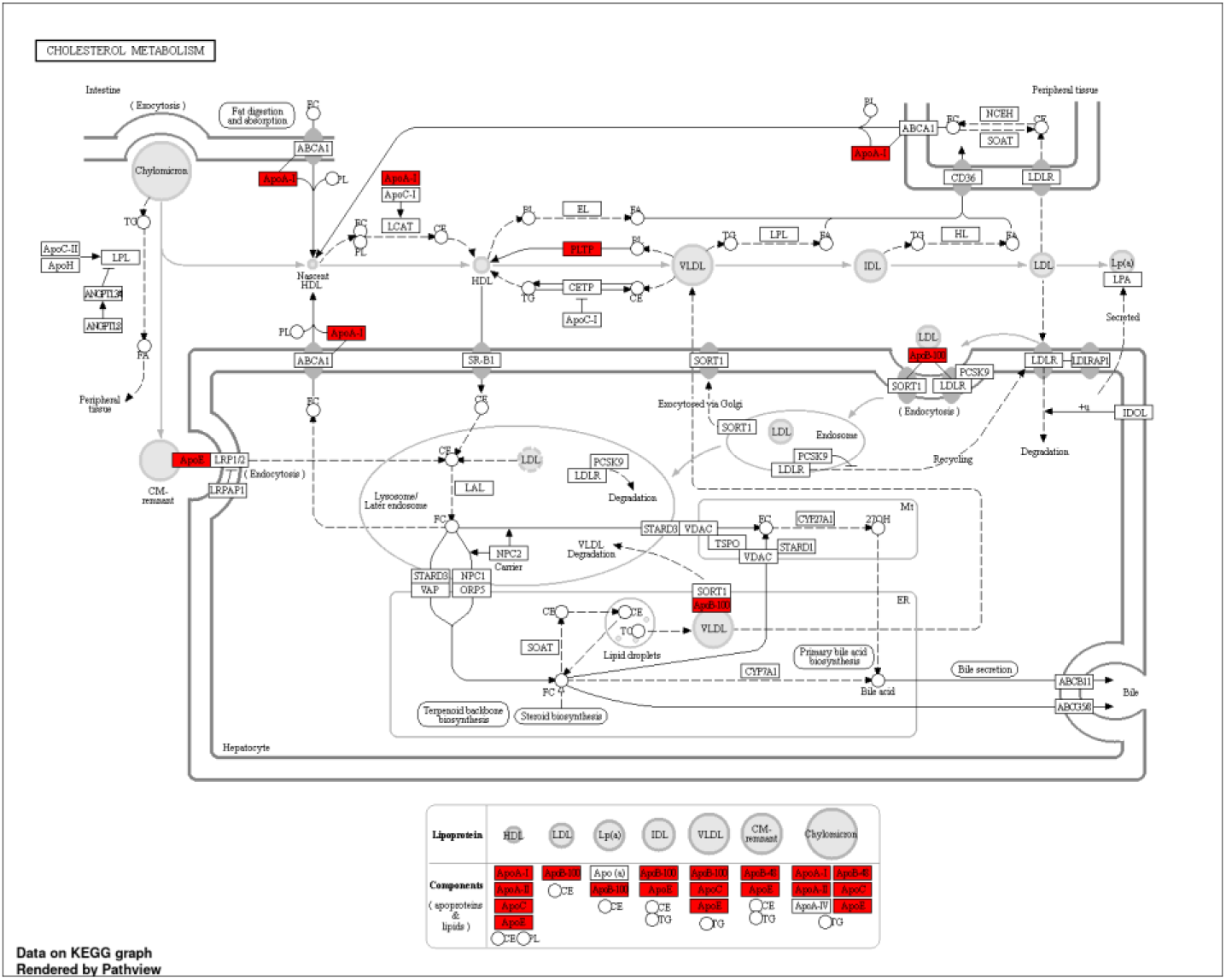
KEGG map of cholesterol metabolism pathway enriched with COT FP SMG.

**Figure S7.**
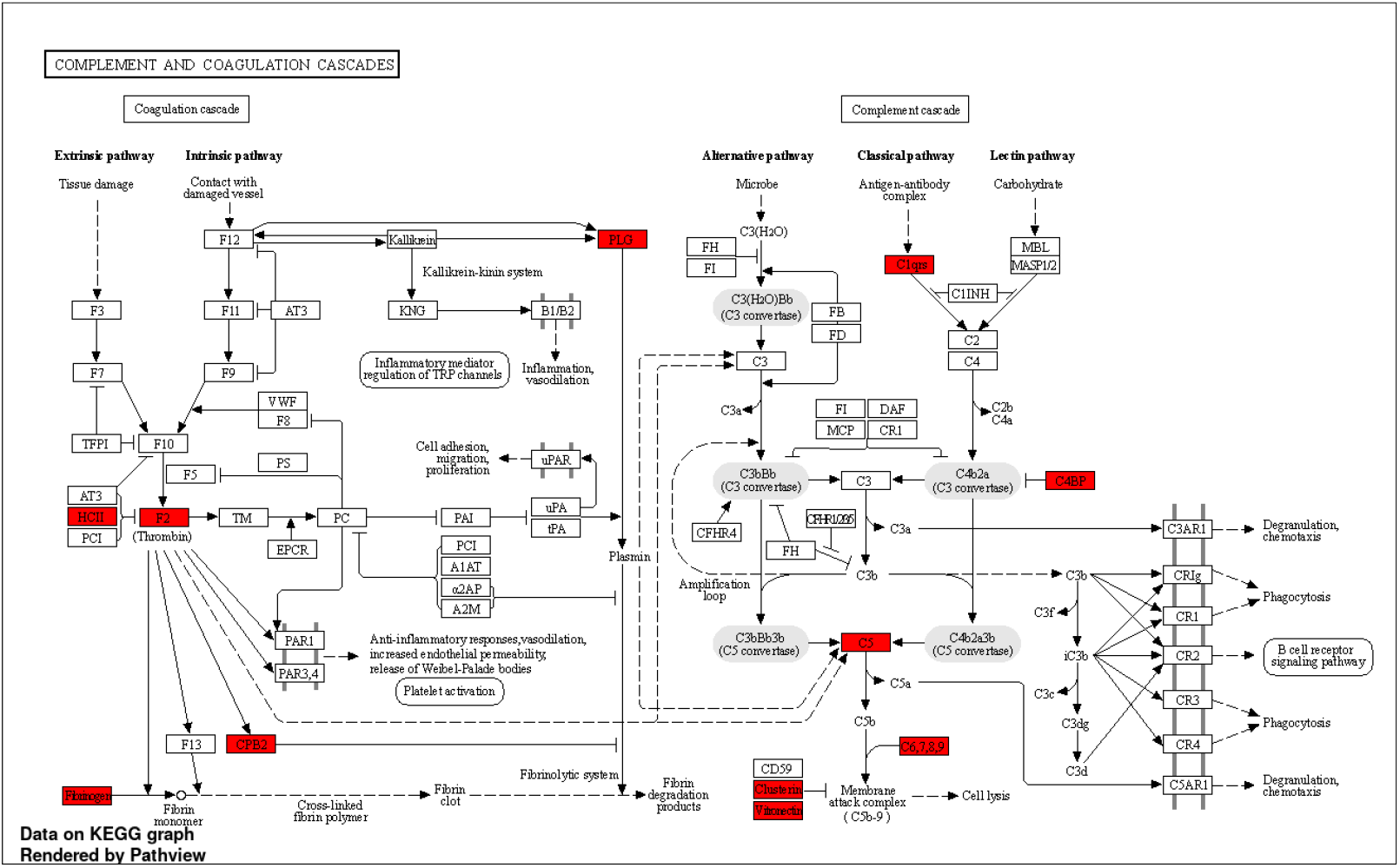
KEGG map of Complement and Coagulation pathway enriched with COT FP SMG.

**Figure S8.**
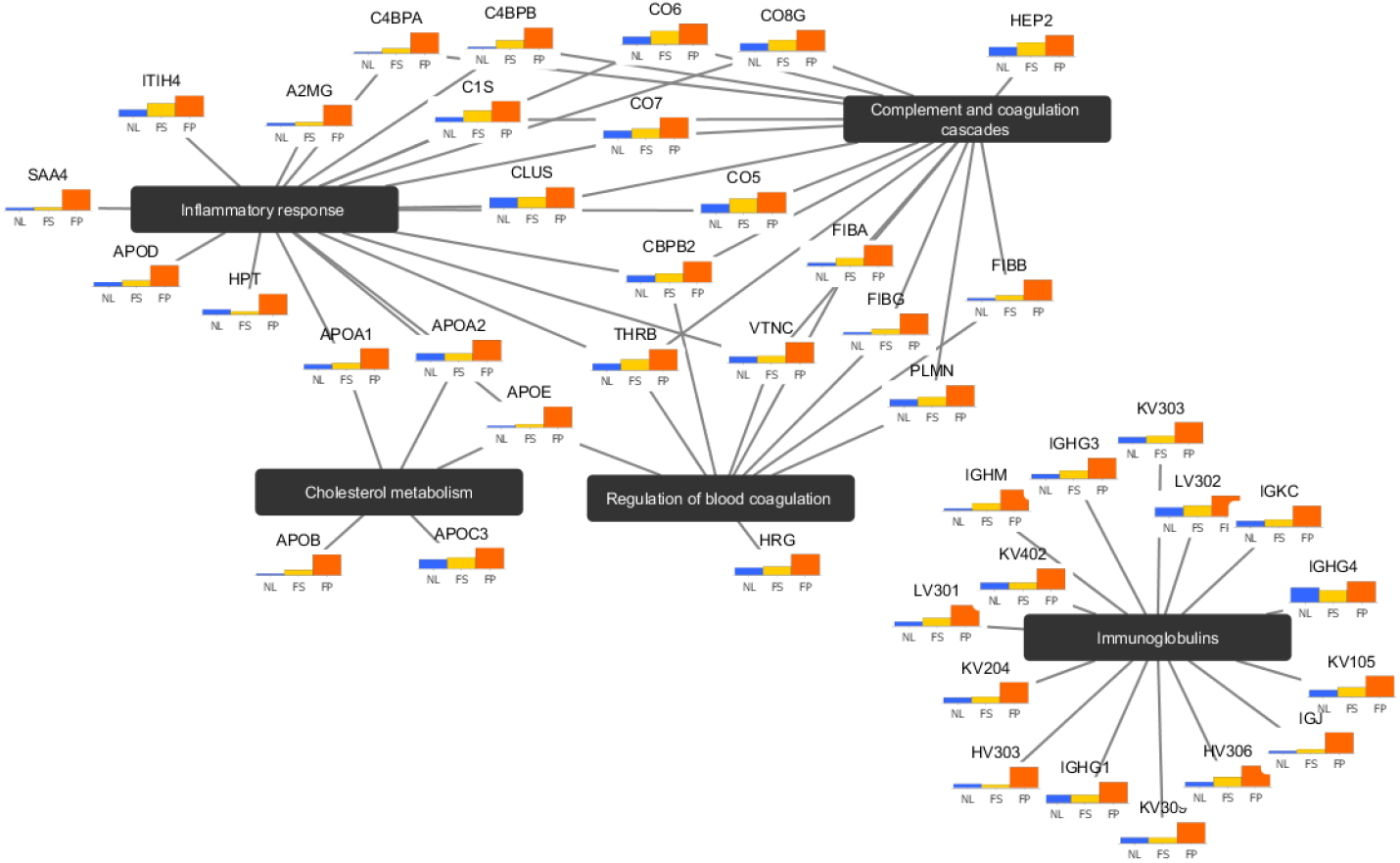
Cytoscape map of the top pathways enriched with COT FP SMG.

**Figure S9.**
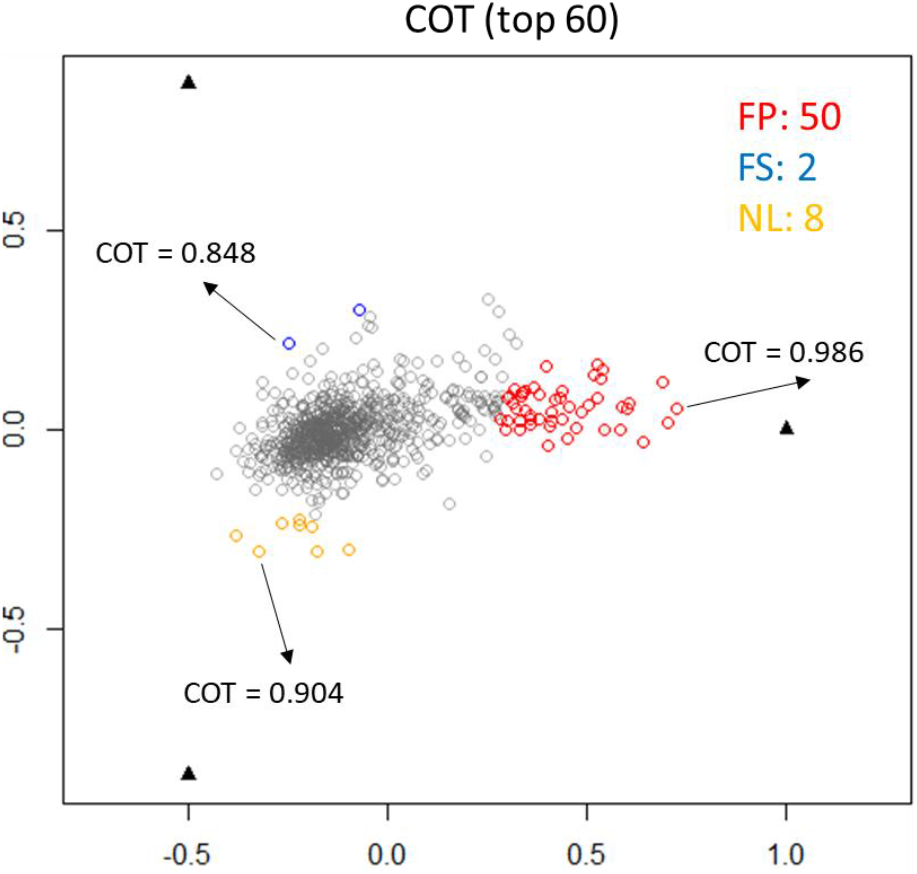
Scatter simplex of color-coded COT top protein SMG detected directly from heterogeneous samples.

## ACKNOWLEDGMENTS

This work has been supported by the National Institutes of Health under Grants HL111362-05A1, HL133932, NS115658-01, and the Department of Defence under Grant W81XWH-18-1-0723 (BC171885P1).

